# Aberrant Neural Entrainment to Word-Level Speech Patterns in Fragile X Syndrome: Evidence for a Statistical Learning Deficit

**DOI:** 10.1101/2025.10.09.681399

**Authors:** Laura Batterink, Yanchen Liu, Grace Westerkamp, Jae Citarella, Peyton Siekierski, Lynxie Voorhees, Lauren E. Ethridge, Elizabeth Smith, Rana Elmaghraby, Craig A. Erickson, Zag ElSayed, Anubhuti Goel, Steve W. Wu, Ernest V. Pedapati

**Affiliations:** Department of Psychology, Western Centre for Brain and Mind, Western Institute for Neuroscience, University of Western Ontario, London, Ontario, Canada; Division of Child and Adolescent Psychiatry, Cincinnati Children’s Hospital Medical Center, Cincinnati, Ohio, USA; Department of Psychiatry, University of Cincinnati College of Medicine, Cincinnati, Ohio, USA; Division of Behavioral Medicine and Clinical Psychology, Cincinnati Children’s Hospital Medical Center, Cincinnati, Ohio, USA; Department of Pediatrics, Section on Developmental and Behavioral Pediatrics, University of Oklahoma Health Sciences Center, Oklahoma City, Oklahoma, USA; Department of Psychology, University of Oklahoma, Norman, Oklahoma, USA; Neuroscience Graduate Program, University of California, Riverside, Riverside, California, USA; Department of Psychology, University of California, Riverside, Riverside, California, USA; Division of Neurology, Cincinnati Children’s Hospital Medical Center, Cincinnati, Ohio, USA; Department of Pediatrics, University of Cincinnati College of Medicine, Cincinnati, Ohio, USA; School of Information Technology, University of Cincinnati, Ohio, USA

**Keywords:** Fragile X syndrome, statistical learning, electroencephalography, EEG, neural entrainment, language development, auditory processing

## Abstract

Fragile X syndrome (FXS), the most common inherited cause of intellectual disability and autism spectrum disorder, causes significant language and cognitive impairments. Statistical learning refers to the ability to extract patterns from sensory input through mere exposure and plays a central role in language acquisition. Surprisingly, statistical learning in FXS has not been explored. Given that children with FXS typically follow a delayed developmental trajectory for language, we hypothesized that they would show impaired statistical learning. To test this hypothesis, we used an EEG measure of neural entrainment to index statistical learning of hidden trisyllabic words within a continuous speech stream in children with FXS (n = 17) and in typically developing controls (n = 31). Children with FXS showed significantly reduced neural entrainment to words compared to controls, particularly in the superior temporal gyrus and transverse temporal gyrus (primary auditory cortex), providing evidence of statistical learning impairment. Notably, syllable-level entrainment was preserved or even enhanced in FXS, indicating that word-level deficits cannot be attributed to general auditory processing impairments. In addition, while typically developing controls showed an increase in word-level entrainment over the course of learning, children with FXS failed to show a similar increase over time. Taken together, this pattern of results demonstrates that children with FXS can process rapid, lower-order acoustic structure but struggle to integrate these syllables into longer, chunk-like word representations. Overall, these findings suggest that statistical learning is impaired in FXS, and also suggest neural entrainment to statistical structure as a potential therapeutic target.

## Introduction

Statistical learning is the ability to extract patterns from sensory input through mere exposure, without instruction, reinforcement, or feedback (Aslin, 2017; Saffran, Aslin, et al., 1996). In the seminal demonstration of statistical learning, 8-month-infants learned the underlying statistical structure of a continuous stream of repeating trisyllabic nonsense words (e.g. “tupiro”) after only 2 minutes of passive listening (Saffran, Aslin, et al., 1996). Subsequent work has confirmed that statistical learning is a powerful learning mechanism that is present in infants, children and adults (Moreau et al., 2022; Raviv & Arnon, 2018; Saffran et al., 1997; Saffran, Newport, et al., 1996) and that it supports widespread cognitive abilities, including visual perception (Fiser & Aslin, 2001), motor learning (Theeuwes et al., 2024), social learning (Nencheva et al., 2025), and especially language acquisition (Saffran & Kirkham, 2018).

Because statistical learning is foundational in language acquisition and other aspects of cognition (e.g., Kidd, 2012; Romberg & Saffran, 2010; Sherman et al., 2020), understanding statistical learning in neurodevelopmental disorders may provide new mechanistic insights for the altered patterns of learning and language outcomes associated with these disabilities (Saffran & Kirkham, 2018). However, to date there are mixed findings regarding statistical learning abilities in NDDs as a whole. For example, children with Developmental Language Disorder (formerly known as Specific Language Impairment) seem to consistently show poorer performance on statistical learning tasks compared to typically developing children (Evans et al., 2009; Haebig et al., 2017; Obeid et al., 2016). In contrast, for autism spectrum disorder (ASD), there is behavioral evidence both for (Hu et al., 2024; Jones et al., 2018) and against statistical learning impairments (Brown et al., 2010; Haebig et al., 2017; Mayo & Eigsti, 2012; Obeid et al., 2016). However, as has been previously pointed out (Saffran & Kirkham, 2018), behavioural evidence pertaining to statistical learning in NDDs is limited to relatively high functioning children who are able to successfully perform explicit behavioral tests (e.g., making forced-choice judgements about which syllable sequences are more familiar). Capturing statistical learning with neural measures allows for inclusion of a wider range of children, including those with more severe deficits who cannot perform behavioral tasks, and may lead to different conclusions. Supporting this idea, several studies to date that have used neural measures of statistical learning indicate reduced learning abilities in children with ASD (Jeste et al., 2015; Scott-Van Zeeland et al., 2010; Wagley et al., 2020), in contrast to the mixed behavioral findings, though more data is needed.

An additional measure of statistical learning that has not been previously used in the investigation of NDDs is neural entrainment (Obleser & Kayser, 2019), which arises through the synchronization of brain oscillations to the frequency of repeating patterns. By presenting speech patterns at a fixed rhythm, neural entrainment can be leveraged to provide an objective, real-time measure of statistical learning (Batterink & Paller, 2017; see Figure 1), especially useful in children with NDDs who are unable to perform behavioral tasks. In healthy infants, children and adults, entrainment at the frequency of repeating regularities reflects the perceptual binding of individual stimuli into integrated units (e.g. syllables to words), and correlates with subsequent performance on behavioural measures of learning (Batterink, 2020; Batterink & Paller, 2017, 2019; Choi et al., 2020; Moreau et al., 2022). In addition, neural entrainment to syllable occurs without requiring focused attention to the speech stream (Batterink & Paller, 2019) and thus may provide an index of statistical learning that is relatively less influenced by other cognitive factors (e.g., attention, executive control).

**Figure 1.**
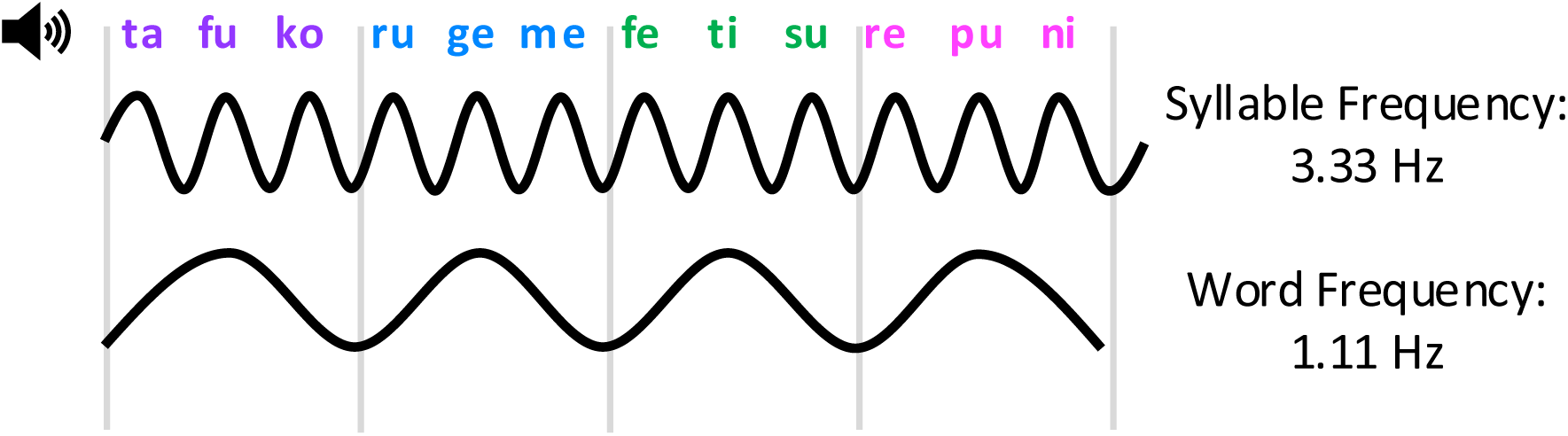
Neural entrainment statistical learning paradigm. Syllables are presented at 3.33 Hz, forming hidden trisyllabic words presented at 1.11 Hz. Neural entrainment at 1.11 Hz, as measured by intertrial phase coherence (ITC), serves as an online neural index of statistical learning. Entrainment at 3.33 Hz reflects sensory tracking of the auditory signal.

Fragile X Syndrome (FXS) provides a unique opportunity to investigate potential statistical learning deficits in NDDs because the underlying biology of this single-gene deficit is very well-characterized, in contrast to other NDDs such as idiopathic autism. FXS results from hypermethylation of the FMR1 gene, leading to loss of Fragile X Messenger Ribonucleoprotein, an RNA-binding protein that regulates activity-dependent translation at synapses. This protein normally acts as a translational repressor for hundreds of mRNAs critical for synaptic plasticity, and its absence results in excessive protein synthesis, disrupted synaptic pruning, and altered excitatory-inhibitory balance. These molecular disruptions manifest clinically as intellectual disability, language delays, and autism symptoms in approximately 60% of males and 20% of females with FXS (Baker et al., 2019; Hagerman et al., 2017). Language development is particularly affected, with pronounced delays and deficits in receptive language, vocabularly, morphology and syntax, conversational discourse, and processing of rapidly changing auditory information (Finestack et al., 2009; Hoffmann, 2022).

Despite extensive documentation of language impairments in FXS, no studies have directly examined statistical learning in this population. This gap is particularly striking given that: (1) statistical learning underlies many linguistic abilities affected in FXS, including word segmentation and syntax acquisition (Romberg & Saffran, 2010); (2) statistical learning can be objectively measured through neural entrainment, requiring nothing more than passive listening on the part of the participant and thus circumventing the behavioral assessment challenges in intellectually disabled populations (Batterink & Paller, 2017, 2019; Batterink & Zhang, 2022; Choi et al., 2020; Moreau et al., 2022). Furthermore, as previously alluded to, the single-gene basis of FXS offers a clear path to understanding biological mechanisms, setting the stage for future work in this area. In the current study, using a direct, neural entrainment measure of statistical learning, we test the hypothesis that children with FXS will show reduced neural entrainment (measured via inter-trial phase coherence) to repeating patterns in auditory streams compared to age-matched typically developing controls, reflecting a deficit in the online computations underlying statistical learning. By documenting statistical learning deficits in FXS, this work provides the first direct evidence of impaired implicit learning mechanisms in this population and establishes a foundation for future mechanistic studies and targeted interventions.

## Methods and Materials

### Ethical Considerations

All procedures were approved by the Cincinnati Children’s Hospital Institutional Review Board (IRB #2015–8425). Written informed consent was obtained from parents or legal guardians, and children provided age-appropriate assent when possible. For younger children or those with intellectual disabilities, willingness to participate was carefully assessed.

### Demographic Information

Participants included individuals with FXS with full mutation confirmed by PCR/Southern blot and typically developing controls (TDC; average IQ 85–115, no neuropsychiatric family history). Behavioral and medication regimens were stable for ≥30 days (≥60 days for non-stimulant psychotropics). Exclusion criteria were unstable seizure disorder, significant medical or neurological illness, unresolved acute illness, peripheral hearing loss, use of GABAergic or glutamatergic modulators, or inability to comply with study procedures.

A total of 17 children with Fragile X syndrome and 31 typical developing children contributed data to the study (please see Table 1 for demographic and IQ scores for each group). Two additional typically developing controls were recruited but later excluded from the final dataset due to poor data quality (n = 1) and a technical problem with the EEG file that precluded data analysis (n = 1).

**Table 1.** Demographic and IQ Data.

| Characteristic | FXS, N = 17<sup>1</sup> | TDC, N = 31<sup>1</sup> | p-value<sup>2</sup> |
| --- | --- | --- | --- |
| Sex | 4 F (24%), 13 M (76%) | 17 F (55%), 14 M (45%) | 0.037 |
| Age (Years) | 9.4 ± 3.0 (5.2 – 14.7) | 9.3 ± 3.0 (5.1 – 15.5) | >0.9 |
| Full Scale Deviation IQ | 50.8±18.5 | 104.8±7.5 | <0.001 |
| Nonverbal Deviation IQ | 51.4±17.403 | 105.3±8.5 | <0.001 |
| Verbal Deviation IQ | 50.1±20.5 | 104.9±9.5 | <0.001 |

<sup>1</sup>n (%); Mean ± SD
<sup>2</sup>Pearson's Chi-squared test; Welch Two Sample t-test

### Stimuli

Syllables contributing to the stream were the same as those used by Batterink and Paller (2019). Syllables were recorded by a male native English speaker using neutral intonation and no co-articulation between syllables. Each syllable was saved within a separate file, with the beginning of each sound file coinciding with the precise onset of the syllable. Syllables were concatenated to create 4 nonsense “words” (“tafuko”, “rugeme”, “repuni”, “fetisu”). To create the continuous speech stream, syllables were presented at a rate of 300 ms per syllable (3.33 Hz) in a predefined pseudorandom order, with the constraint that the same word did not repeat consecutively. Thus, trisyllabic words were presented at a rate of 1.11 Hz. The speech stream contained a total of 2400 syllables (800 words), with each word presented 200 times each (12 min total duration).

### Procedure

After electrode setup for EEG recording, participants were seated in a sound-attenuated booth at a comfortable viewing distance from a computer monitor. The continuous speech stream was presented with Sony MDR-V150 headphones at 65 dB while participants’ EEG was recorded.

To facilitate cooperation, participants viewed a silent video of their choice, presented without words or captions, while the speech stream was presented.

### EEG Recording and Preprocessing

#### Scalp Level Preprocessing

EEG data were recorded at a 1000 Hz sampling rate with an EGI NetAmp 400 with a 128-channel HydroCel electrode net (Magstim/EGI, Eugene, OR). EEG preprocessing and analysis followed the basic approach used in our prior study that used the same recording system (Choi et al., 2020). First, a 60-Hz notch filter and a band-pass Butterworth filter from 0.5 to 20 Hz were applied to the raw data. Next, we performed artifact correction using the Artifact Blocking (AB) (Mourad et al., 2007) algorithm to attenuate artifacts (e.g. eye blinks, eye movements, and body movements). As recommended by Fujioka and colleagues we applied the algorithm after removing the outer ring of channels from further analysis, and we set the threshold value within the artifact-block algorithm at ±50 μV. Visual inspection of the individual data confirmed that the artifact-block algorithm successfully corrected artifacts and reduced noise in the data. The artifact-corrected data were then re-referenced to a common average. Nonoverlapping epochs of 9.0 s, time-locked to the onset of every 10th word and corresponding to a duration of 10 words, were then extracted, yielding 78 epochs per participant.

#### Source Localization

To conduct parallel analyses at the source level, the cleaned scalp EEG data was used for source-modeling with the fsaverage template brain in MNE 1.9 (Gramfort et al., 2010). Individual MRIs and electrode digitization were not available; therefore, source localization was employed primarily as an optimal spatial filter rather than for precise anatomical mapping. This approach was particularly important for separating activity in temporal and frontal regions, given the orientation of Heschl’s gyrus and its proximity to frontal sources. The forward solution was computed with a precomputed BEM and ico-5 source space (minimum source–sensor distance: 5 mm), and the inverse operator was estimated with λ² = 1/9 and fixed-orientation dipoles. Source time courses (STCs) were parcellated into 68 cortical nodes using the Desikan–Killiany atlas (Desikan et al., 2006) and obtained for each epoch.

### Computation of Intertrial Phase Coherence and Statistical Analysis

For each participant, we quantified neural entrainment by measuring inter-trial coherence (ITC) across the nonoverlapping epochs. ITC is a measure of event-related phase-locking or phase synchronization across trials, which ranges from 0 to 1, with 0 indicating purely non-phase-locked (i.e., random) activity at a given frequency band, and 1 indicating strictly phase-locked activity. Higher ITC values indicate more consistency in the phase of the signal across individual trials (in our case, epochs time-locked to word onsets). To the extent that statistical learning occurs, we expected to observe higher phase-locking at the triplet frequency, reflecting greater neural entrainment to the hidden word structure. The Fast Fourier transform was applied to each epoch to decompose the signal into its frequency components, and ITC was then computed across all epochs across frequencies, including each frequency of interest (word and syllable). For scalp level analyses, this procedure was carried out for all electrode channels. For source level analyses, this procedure was conducted for all 68 nodes.

To assess statistical significance of ITC values, for each channel (scalp level analysis) or node (source level analysis), we converted raw ITC values to z-scores of ITC (zITC), thereby normalizing each individual’s ITC values against their own null distribution of non-entrained activity. To accomplish this, we generated 100 surrogate datasets for each participant. To create each surrogate dataset, each epoch onset was individually shuffled by a different random value between –900 ms and 900 ms from the actual epoch onsets (following (Batterink & Zhang, 2022; Herrera-Chaves et al., n.d.; Moreau et al., 2022). This procedure eliminates the alignment between the EEG signal and the auditory speech stream, while preserving the general features of the data and each epoch’s general timing within the experimental task. ITC was then computed for each of the 100 surrogate datasets, resulting in a null distribution of ITC values (et each channel/node). Next, zITC for each electrode was computed using the standard z-score formula (i.e., subtracting the mean of the surrogate ITCs from the observed ITC, and dividing this value by the standard deviation of the surrogate ITCs).

#### Scalp-Level Statistical Analysis

Mean zITC values were averaged across electrodes to produce an average entrainment value for each participant, at each frequency of interest (Word, Syllable). We extracted zITC values for all channels because we observed relatively widespread ITC effects across the scalp at both frequencies of interest (see Figure 2). To test whether neural entrainment at each frequency of interest differed between groups, we conducted two separate univariate ANOVA with Group (TDC, FXS) as a between-subject factor and mean zITC_word_ (or zITC_syllable_) across channels as the dependent measure. We additionally tested whether each group showed significant neural entrainment at each frequency of interest by conducting a one-sample t-test against 0, two-tailed.

**Figure 2.**
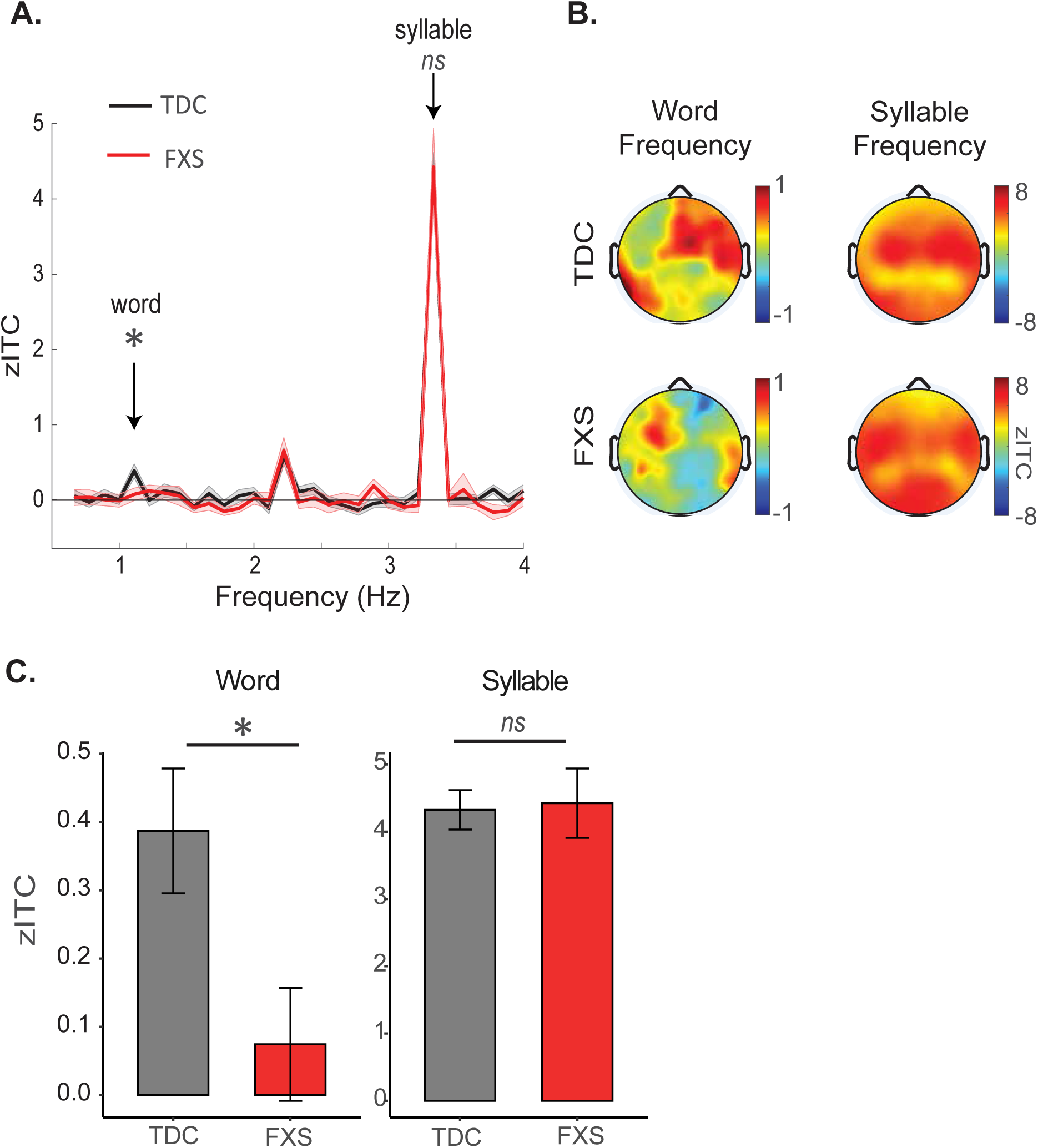
EEG results from scalp-level analyses. A) Z-scored ITC values (zITC) as a function of group (TDC in black; FXS in red) and frequency. Children with FXS show a significant reduction in neural entrainment at the word frequency (1.11 Hz); no group differences are observed at the syllable frequency (3.33 Hz). EEG data is averaged across scalp electrodes. Shaded error bars indicate standard error of the mean. B) Topographical plots showing distribution of zITC across the scalp, as a function of Group and Frequency. C) zITC values at our two frequencies of interest, summarized from data shown in panel A.

#### Source-Level Statistical Analysis

We examined group differences in source-level entrainment using linear mixed-effects models (LMMs) that accounted for the nesting of nodes within cortical regions. Region and Hemisphere were both sum coded, and Group was treatment coded with the TDC group serving as the reference group.

### Word-Level Entrainment

For word-level responses (ZITC_word_), the primary model (WordModel1) included Group (TDC, FXS), Cortical Region (prefrontal, frontal, temporal, central, parietal, lingual, occipital), Hemisphere (left, right), and their interactions, with random intercepts for participant and node:

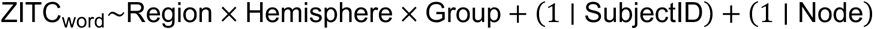

If a significant Group × Region interaction was found, this motivated a refined **node-level model** (WordModel2):

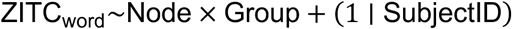

Follow-up comparisons were conducted using estimated marginal means (emmeans), testing group differences at the node level without multiple-comparison adjustment.

### Syllable-Level Entrainment

For syllable-level responses (ZITC_syllable_), we used an analogous region-level model:

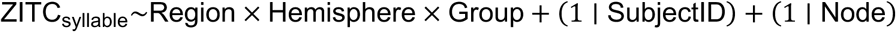

#### Word Precision Index

Within bilateral transverse temporal gyrus nodes (identified as key sites in our main analyses), we further quantified the precision of phase locking at the word frequency using a novel measure we term the *Word Precision Index (WPI)*. This index reflects the selectivity of neural entrainment to words relative to nearby frequencies and was computed as:

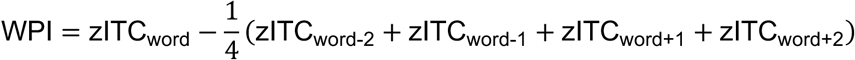

where zITC_word_ is the entrainment at the target word frequency and zITC_word±n_ are the four neighboring bins. Higher values indicate more precision (accurate) phase locking to the component words in the stream. We tested for group differences in the word precision index using a two-sample t-test.

### Time Course of Neural Entrainment Over Learning

In our prior studies in healthy infants, children and adults, we have observed an increase in neural entrainment at the word frequency over the exposure period, reflecting the progression of statistical learning over time (Batterink, 2020; Batterink & Paller, 2017, 2019; Choi et al., 2020; Moreau et al., 2022). These increases appear most consistently within the first half (∼6 minutes) of exposure to the artificial language (Batterink & Paller, 2019; Choi et al., 2020; Moreau et al., 2022), after which entrainment values fluctuate around an asymptotic value rather than continuing to increase (Batterink & Paller, 2019). Therefore, to quantify the progression of learning over time in the current study, we focused on examining neural entrainment values at our frequencies of interest within the first half of exposure.

Since ITC is undefined at the single trial level, we used a jackknifing approach (Richter et al., 2015; Waschke et al., 2019) to estimate ITC values at the single epoch level at our two frequencies of interest (word and syllable). In this procedure, ITC is initially computed across all epochs (as in our main analysis). Next, ITC is recomputed on all epochs but the first one, which represents one leave-one-out jackknife replication. This procedure is repeated for each epoch (systematically leaving out one epoch each time), producing a total of N jackknife replications, where N is the total number of epochs. These jackknifed ITC values represent the values that the overall ITC would have had without that specific trial. We normalized these jackknifed ITC values at each frequency of interest by subtracting the average of the jackknifed ITC estimates for all other frequency bins under 5 Hz, excluding the word and syllable frequencies and their harmonics (i.e., frequencies spanning from 0.66 to 5.0 Hz, excluding frequency bins 1.11, 2.22, 3.33 and 4.44 Hz). For ease of interpretability, we then converted each of these normalized jackknifed ITC values to ITC pseudovalues (using the formula N*ITC_all_– (N-1)*Jackknifed_ITC). The ITC pseudovalue for a given epoch represents the contribution of that epoch to the overall ITC. That is, the higher the pseudovalue, the more a given trial contributes to increasing the overall ITC, as its phase at a given frequency resembles the mean phase at that frequency across trials. Finally, because ITC estimates at the epoch level are inherently very noisy, we smoothed the pseudovalues using a moving average with a span of 5 datapoints (i.e., each nth pseudovalue was averaged with neighbours n-2, n-1, n + 1 and n+2) to reduce the influence of outliers in our statistical analysis, which excludes the first two and final two datapoints from further analysis. We hypothesized that these trial-level ITC estimates (pseudovalues) should increase over time in the TDC group, tracking the process of statistical word learning, and that this increase may be less robust in the FXS group, reflecting impairments in the statistical learning process.

We statistically tested whether neural entrainment at our frequencies of interest changed over time by modeling the smoothed ITC pseudovalues at each frequency of interest using a linear mixed model with Epoch Number, Group and their interaction as fixed factors, and participant as a random intercept (Time Course Model: [ITC_Pseudovalue_word_ ∼ EpochNumber*Group + (1 | SubjectID). Group was again treatment coded with the TDC group serving as the reference group. Given the results from our main analysis, we conducted this entire procedure at the scalp level (averaging pseudovalues across all electrodes) and again at the source level, within bilateral transverse temporal gyrus nodes, where neural entrainment effects were maximal.

## Results

### Scalp-Level Analysis

Consistent with our hypothesis, children with FXS showed significantly reduced neural entrainment at the word frequency compared to the TDC group (zITC_word_ Group effect: (*F* (1,46) = 5.11, *p* = 0.028; Figure 2). Further, while TDC participants showed highly significant word-level entrainment as a group (ZITC_word_ > 0: t(30) = 4.23, p < 0.001), word entrainment in the FXS group was not significant (ZITC_word_ > 0: t(16) = 0.90, p = 0.38). In contrast, syllable-level entrainment did not differ significantly between the two groups (zITC_syllable_ Group effect: *F* (1,46) = 0.033, *p* = 0.86), and both groups showed highly significant entrainment at the syllable frequency (both p values < 0.001). Overall, these results suggest a highly specific and aberrant pattern of reduced neural entrainment to words in the FXS group.

### Source-Level Analysis

#### Word Entrainment

The TDC group showed highly robust neural entrainment to words, which was significant across the brain but particularly strong in the temporal cortex (WordModel1: mean parameter estimate: 0.27, SEM = 0.060, p < 0.001; temporal cortex estimate: 0.40, SEM = 0.056, p < 0.001; see Figure 3 and Figure 4; Table S1). No significant main effects of hemisphere or interactions with hemisphere were observed, indicating that entrainment was similarly robust across both hemispheres.

**Figure 3.**
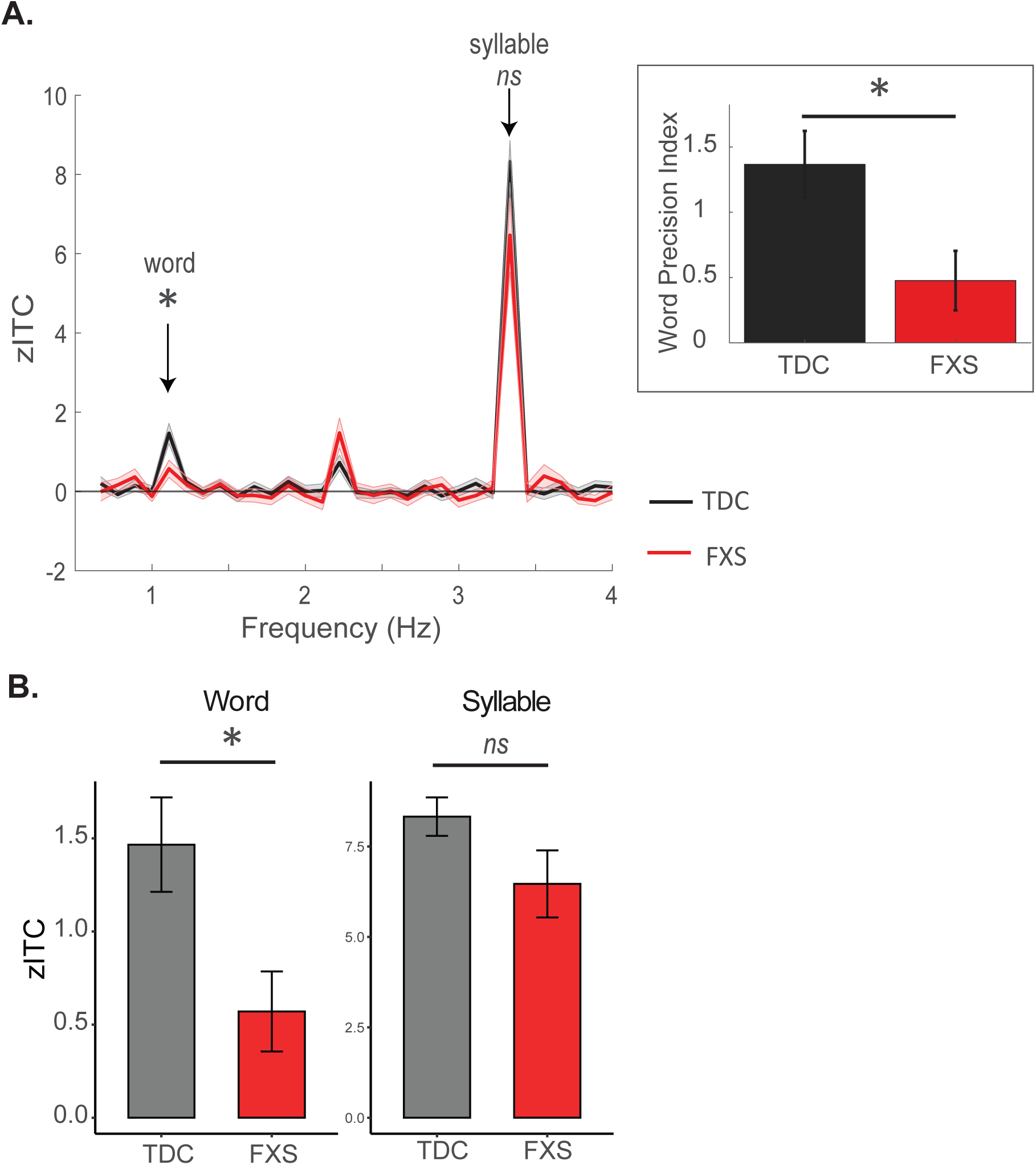
EEG results from source-level analyses; all data shown is drawn from bilateral transverse temporal gyrus (primary auditory cortex), where maximal neural entrainment was observed. A) Z-scored ITC values (zITC) as a function of group and frequency. Children with FXS show a significant reduction in neural entrainment at the word frequency (1.11 Hz). Inset – entrainment values at the word and neighbouring frequencies are used to compute the word precision index. Children with FXS show significantly reduced precision in word tracking. B) zITC values at our two frequencies of interest, summarized from data shown in panel A.

**Figure 4.**
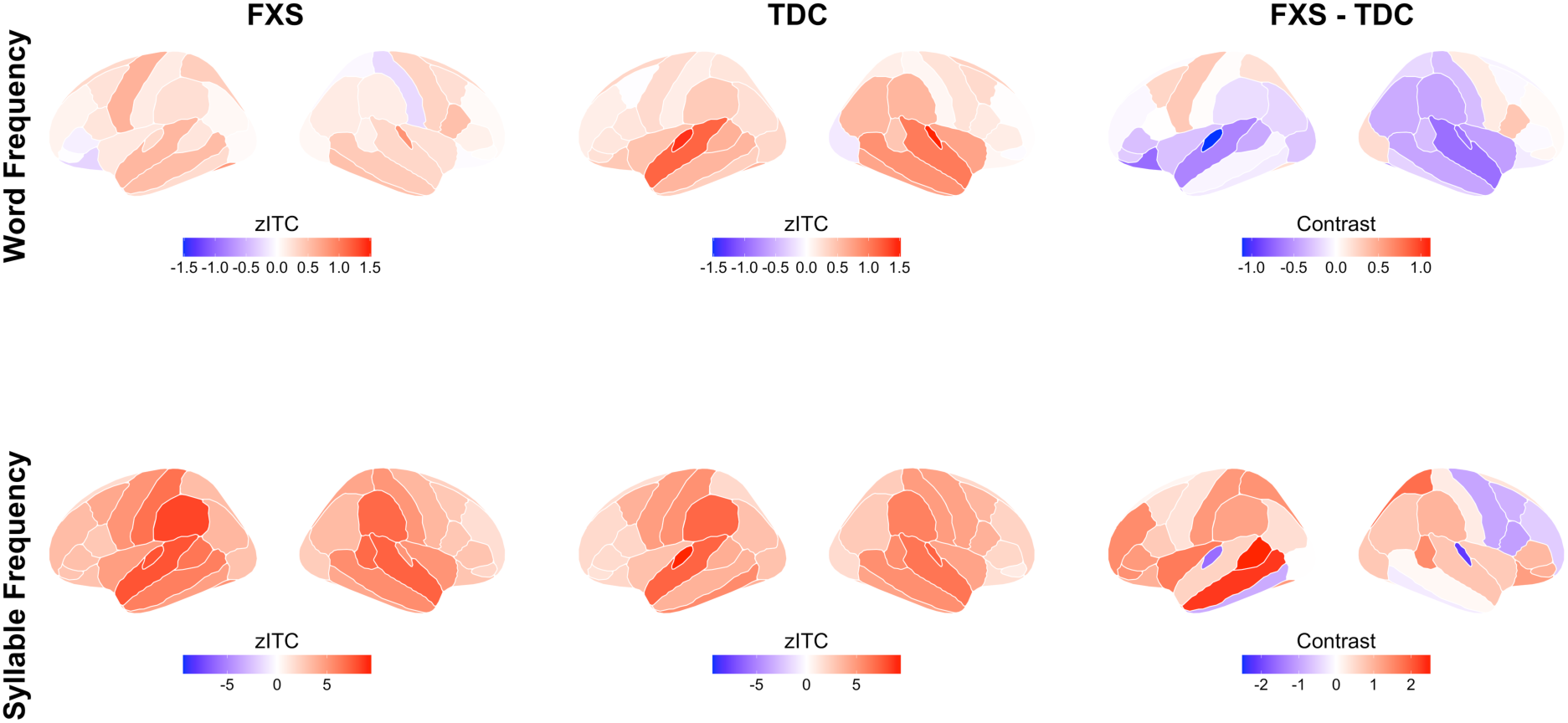
Neural entrainment to word frequency is reduced in FXS. Lateral views of cortical source-localized z-scores of ITC (zITC) for word frequency (top row) and syllable frequency (bottom row) processing for left and right hemispheres. Left column shows FXS group averages, middle column shows TDC group averages, and right column shows group numerical differences (FXS – TDC). Color scales represent zITC values, where positive values (red) indicate greater phase-locking to the respective frequency compared to surrogate data, and negative values (blue) indicate reduced phase-locking. Source localization was performed using the Desikan-Killiany atlas with 68 cortical nodes.

Of relevance to our main hypothesis that children with Fragile X would exhibit reduced neural evidence of statistical learning compared to healthy controls, our initial model revealed a robust Group x Temporal Region interaction (Temporal Region x Group: –0.25, SEM = 0.078, p= 0.002). A follow-up contrast confirmed that the FXS group showed significantly lower word entrainment than the TDC group within temporal cortex (estimate = 0.33, SE = 0.12, p = 0.004); no other regions showed significant group differences (*p* > 0.1). A second model at the Node level demonstrated that FXS children showed significantly reduced entrainment within bilateral superior temporal gyrus (TDC – FXS estimate: 0.64, SE = 0.26, z ratio = 2.49, p = 0.013) and within bilateral transverse temporal gyrus (TDC – FXS estimate: 0.90, SE = 0.26, z ratio = 3.49, p = 0.005). In addition, FXS children also showed significantly reduced precision in neural entrainment to the word frequency within the transverse temporal gyrus node, as reflected by the word precision index, relative to TDC children (t (46) = 2.31, p = 0.025; Figure 3A, inset). This result reflects the FXS group’s “blurred” entrainment profile of above-zero entrainment in frequency bins adjacent to the word frequency, indicative of imprecise or inaccurate phase-locking. In summary, relative to typically developing children, children with Fragile X show significantly reduced entrainment to words in both superior temporal and transverse temporal gyrus.

#### Syllable Entrainment

Within the TDC group, highly robust neural entrainment effects at the syllable frequency were observed (estimate = 2.98, SE = 0.29, p < 0.001). These entrainment effects were maximal over temporal and central cortex (Temporal Region: estimate = 1.71, SEM = 0.32, p < 0.001; Central Region: estimate = 1.53, SEM = 0.51, p = 0.004). Again, no significant main effects of hemisphere or interactions with hemisphere were observed, indicating that entrainment was similarly robust across hemispheres. No significant main effect of group was observed (p = 0.15), although the Fragile X group was associated with numerically higher zITC estimates. In addition, in contrast to the general pattern of reduction observed for word entrainment, Fragile X children showed significantly stronger syllable entrainment in the left hemisphere and in parietal cortex compared to typically developing children (Group x Hemisphere: estimate = 0.31, SEM = 0.010, p = 0.002); Group x Parietal Region: estimate = 0.053, SEM =0.025, p = 0.035). This result suggests that the diminished word-level entrainment in Fragile X cannot be attributed to a general reduction in neural responsiveness to auditory stimuli.

### Time Course of Neural Entrainment

#### Scalp Level (All Scalp Electrodes)

Our time course model revealed that word entrainment in the TDC group increased over the first half of exposure as hypothesized, as reflected by a significant epoch number effect (t (1726) = 2.90, p = 0.004). There was no interaction between epoch number and group (t (1726) = –0.94, p = 0.35), suggesting that the two groups showed a similar increase in neural entrainment over the first half of exposure. However, follow-up contrasts using *emtrends* against 0 indicated that the increase in word entrainment over time within the FXS group was not significant (FXS: slope = 0.0004, SE = 0.0005, t (1726) = 0.98, *p* = 0.33; TDC: slope = 0.001, SE = 0.0003, t (1726) = 2.9, p = 0.004).

The same analysis computed for syllable-level entrainment showed a reduction in syllable entrainment over time within the TDC group (epoch number estimate: –0.00074, SE = 0.00036, t (1726) = –2.09, p = 0.037). This decrease over time was not significantly different between the two groups (t (1726) = 0.32, p = 0.75).

#### Source Level (Transverse Temporal Gyrus)

Resembling the scalp-level analysis, within the TDC group, word entrainment increased over time (epoch number: t(1726) = 3.18, p = 0.0015). There was a marginally significant interaction between epoch number and group (t (1726) = – 1.74, p = 0.081), indicating that children with Fragile X showed a somewhat smaller increase in word entrainment over time relative to the TDC group. Follow-up contrasts with *emtrends* confirmed that whereas word entrainment significantly increased over time within the TDC group, word entrainment within the FXS group did not increase significantly (FXS: slope = 0.00016, SE = 0.00085, t (1726) = 0.19, p = 0.85; TDC: slope = 0.002, SE = 0.00063, t (1726) = 3.18, p = 0.0015).

The same analysis computed on syllable entrainment revealed no significant change over time within the TDC group (Epoch Number: (t (1726) = 0.25, p = 0.80). Compared to the TDC group, the Fragile X group showed initially smaller entrainment at baseline (Group: t(55) = –2.7, p = 0.009), coupled with a significantly greater increase over time (Epoch Number x Group: t(1726) = 2.49, p = 0.013). Follow-up contrasts with *emtrends* indicated that syllable entrainment significantly *increased* over time within the FXS group (slope = 0.002, SE = 0.0007, t (1726) = 3.29, p = 0.001), in contrast to the stable pattern observed within the TDC group (slope = 0.0001, SE = 0.0005, t (1726) = 0.25, p = 0.80).

## Discussion

Supporting our hypothesis that children with FXS would show impairments in the computations underlying statistical learning, we have identified a selective impairment in word-rate neural entrainment (∼1.1 Hz) and reduced learning-related slope over time in children with FXS. The reduction in statistical learning was robust at the scalp and source levels, with peak effect localized to the primary auditory cortex and adjacent speech-sensitive cortical regions (represented by the transverse temporal gyri and superior temporal gyrus [STG]). The observed dissociation between impaired word-level entrainment alongside potentially heightened syllable-rate entrainment (∼3.3 Hz) represents a particularly striking and meaningful distinction, implicating a deficit in longer-timescale integration, rather than a generalized auditory or attentional limitation in FXS. These finding also highlight word-level entrainment as a candidate biomarker for language-related vulnerabilities and treatment monitoring in FXS.

Neural oscillation models of speech processing show that cortical rhythms parse speech across nested timescales, with low frequency rhythms jointly supporting segmentation from syllables to words (Ding et al., 2016; Gross et al., 2013; Poeppel & Assaneo, 2020). This hierarchical organization reflects cross-frequency coupling mechanisms where slower oscillations modulate faster ones, enabling the brain to package incoming speech information into units of appropriate temporal granularity (Giraud & Poeppel, 2012). Our word versus syllable dissociation maps onto this temporal hierarchy, suggesting that in FXS, the coordinated multi-frequency dynamics needed for constructing hierarchical linguistic representations, particularly the integration processes operating at word-level timescales (∼1-2 Hz), are selectively compromised, while syllable-level processing remains intact (Zioga et al., 2023).

At a systems-level, loss of FMRP in FXS results in disruption of activity-dependent synaptic regulation (Antoine et al., 2019; Gibson et al., 2008), producing local hyperexcitability, degraded temporal fidelity (X. Wang et al., 2023), and excitation–inhibition imbalance. These cellular perturbations manifest as abnormalities in large-scale oscillatory dynamics. In response to an auditory chirp stimulus, participants with FXS demonstrated reduced gamma-band phase locking, markedly elevated background gamma power, and increases in frontotemporal information flow, suggestive of impaired top-down regulation (Pedapati et al., 2025). Together with resting-state evidence of disrupted alpha–theta coupling (Pedapati et al., 2022; J. Wang et al., 2017), reduced alpha power (Van Der Molen et al., 2014; Van der Molen & Van der Molen, 2013), and altered alpha/gamma connectivity (Schmitt et al., 2022), these findings converge on a model of local hyperexcitability and impaired cross-frequency coordination, which in turn reduce signal-to-noise ratio and weaken the scaffolding required for integrating information across timescales. This overall framework helps explain our EEG results: word-level entrainment (a slower rhythm requiring stable cross-frequency interactions and top-down support) is reduced, whereas syllable-level entrainment (a faster, locally driven rhythm) remains intact or is even heightened under conditions of hyperexcitability.

It is notable that we found the strongest group differences within STG and transverse temporal gyrus. The finding of robust entrainment within these regions in the healthy control group aligns well with prior work that has implicated both the transverse temporal gyrus (Herrera-Chaves et al., n.d.; McNealy et al., 2006) and surrounding STG (Cunillera et al., 2009; Henin et al., 2021; Herrera-Chaves et al., n.d.; Karuza et al., 2013; McNealy et al., 2006) as regions forming a central hub in auditory-linguistic statistical learning. Transverse temporal gyrus is the first cortical region to receive and process raw auditory input, while the STG is a critical locus for the processing of speech sounds, integrating lower-level, acoustic-phonetic information with higher-order contextual and linguistic information to support speech perception (Bhaya-Grossman & Chang, 2022; Yi et al., 2019). Through recurrent connections, the STG generates context-dependent representations that capture longer temporal sequences and give rise to word representations (Bhaya-Grossman & Chang, 2022). In the case of statistical learning, the STG’s ability to integrate bottom-up, incoming speech (e.g, a word-final syllable such as “ko”) with the prior context (e.g., the two syllables “ta” and “fu” as in the word “tafuko”) may give rise to a unique distributed pattern of neural activity, which may then form the basis for identification of repeated patterns in continuous speech, as reflected by entrainment (Endress, 2024). Circuits within STG may thus provide a core, automatic mechanism for statistical learning, operating without intentionality or top-down control – a hallmark of statistical learning. The finding that entrainment is impaired within STG in FXS converges broadly with other evidence suggesting that the STG is a vulnerable region in FXS. For example, individuals with FXS show age-related decreases in the volume of the STG (Reiss et al., 1994), a pattern not seen in controls, and reduced gray matter volume of the left STG (Sandoval et al., 2018). FXS individuals also exhibit reduced gamma band synchronization to an auditory chirp stimulus in left and right temporal lobes, as revealed through source-localized EEG (Pedapati et al., 2025).

Learning-over-time dynamics also provide mechanistic insight into why individuals with FXS acquire new information more slowly. Word-rate entrainment in controls, as in previous studies, increased with exposure, representing the accumulating knowledge of word-like regularities(Batterink, 2020; Batterink & Paller, 2017, 2019; Choi et al., 2020). In children with FXS, the word learning rate was essentially flat, suggesting reduced capacity to accrue longer timescale learning, and is consistent with broader findings of slower learning rates and reduced adaptation in FXS across multiple domains (Goel et al., 2018; Knox et al., 2012; Schmitt et al., 2023). Overall, the reduced statistical learning abilities in individuals with FXS may contribute to their known deficits across many components of language (Finestack et al., 2009; Hoffmann, 2022).

A strength of the study is that we used a direct neural measure of statistical learning, which allowed for inclusion of more severely affected participants in our FXS sample who would have had difficulty performing behavioral tasks. However, a limitation of the current study is that the FXS sample was underpowered to test possible correlations between our entrainment measure of statistical learning and cognitive outcomes such as IQ, especially given the wide age range of the sample. In healthy adults, both EEG (van der Wulp et al., 2025) and behavioral measures of statistical learning (Siegelman & Frost, 2015) have been found to be largely unrelated to general cognitive abilities, such as IQ, verbal working memory, and rhythmic abilities, suggesting that in normal development, statistical learning operates as a basic mechanism that does not limit higher-level cognition. In contrast, if functioning of the normal statistical learning capacity breaks down or is atypical, as is the case in NDDs such as FXS, statistical learning might relate more strongly to general cognitive outcomes, in line with prior findings that populations outside of typical development show weaker statistical learning (Evans et al., 2009; Gabay et al., 2015; Lammertink et al., 2017; Vandermosten et al., 2019; Zhang et al., 2021). Testing a possible direct link between altered statistical learning and cognitive outcomes will be an important direction for future work.

To conclude, these results indicate aberrant neural entrainment to higher-order regularities in FXS, consistent with a specific deficit in assembling multi-syllabic units from continuous speech, rather than a global impairment of auditory responsiveness. An important distinction in our findings is the preserved syllable-level entrainment at 3.3 Hz versus impaired word-level entrainment at 1.1 Hz in FXS participants. The pattern suggests that children with FXS can synchronize to rapid, lower-order acoustic structure but struggle to form the longer, chunk-like representations characteristic of word segmentations. Similarly, the time-course analysis demonstrated a characteristic build-up of word-locked entrainment in control participants, but no reliable increase in FXS. In a clinical context, there is early evidence that entrainment growth can be enhanced (i.e., through transcranial magnetic stimulation) (Smalle et al., 2022) and thus may represent mechanistically grounded biomarker (Sahin et al., 2019) that could quantify both learning capacity and treatment efficacy in FXS interventions targeting language development.

## References

1. Antoine, M. W., Langberg, T., Schnepel, P., & Feldman, D. E. (2019). Increased Excitation-Inhibition Ratio Stabilizes Synapse and Circuit Excitability in Four Autism Mouse Models. Neuron, 101(4), 648–661.e4. 10.1016/j.neuron.2018.12.026

2. Aslin, R. N. (2017). Statistical learning: A powerful mechanism that operates by mere exposure. Wiley Interdisciplinary Reviews. Cognitive Science, 8(1–2). 10.1002/wcs.1373

3. Baker, E. K., Arpone, M., Aliaga Vera, S., Bretherton, L., Ure, A., Kraan, C. M., Bui, M., Ling, L., Francis, D., Hunter, M. F., Elliott, J., Rogers, C., Field, M. J., Cohen, J., Santa Maria, L., Faundes, V., Curotto, B., Morales, P., Trigo, C., … Godler, D. E. (2019). Intellectual functioning and behavioural features associated with mosaicism in fragile X syndrome. Journal of Neurodevelopmental Disorders, 11(1), 41. 10.1186/s11689-019-9288-7

4. Batterink. (2020). Syllables in Sync Form a Link: Neural Phase-locking Reflects Word Knowledge during Language Learning. Journal of Cognitive Neuroscience, 32(9), 1735–1748. 10.1162/jocn_a_01581

5. Batterink, L. J., & Paller, K. A. (2017). Online neural monitoring of statistical learning. Cortex, 90, 31–45. 10.1016/j.cortex.2017.02.004

6. Batterink, L. J., & Paller, K. A. (2019). Statistical learning of speech regularities can occur outside the focus of attention. Cortex, 115, 56–71. 10.1016/j.cortex.2019.01.013

7. Batterink, L. J., & Zhang, S. (2022). Simple statistical regularities presented during sleep are detected but not retained. Neuropsychologia, 164, 108106. 10.1016/j.neuropsychologia.2021.108106

8. Bhaya-Grossman, I., & Chang, E. F. (2022). Speech Computations of the Human Superior Temporal Gyrus. Annual Review of Psychology, 73(1), 79–102. 10.1146/annurev-psych-022321-035256

9. Brown, J., Aczel, B., Jiménez, L., Kaufman, S. B., & Grant, K. P. (2010). Intact implicit learning in autism spectrum conditions. Quarterly Journal of Experimental Psychology, 63(9), 1789–1812. 10.1080/17470210903536910

10. Choi, D., Batterink, L. J., Black, A. K., Paller, K. A., & Werker, J. F. (2020). Preverbal Infants Discover Statistical Word Patterns at Similar Rates as Adults: Evidence From Neural Entrainment. Psychological Science, 095679762093323. 10.1177/0956797620933237

11. Cunillera, T., Càmara, E., Toro, J. M., Marco-Pallares, J., Sebastián-Galles, N., Ortiz, H., Pujol, J., & Rodríguez-Fornells, A. (2009). Time course and functional neuroanatomy of speech segmentation in adults. NeuroImage, 48(3), 541–553. 10.1016/j.neuroimage.2009.06.069

12. Desikan, R. S., Ségonne, F., Fischl, B., Quinn, B. T., Dickerson, B. C., Blacker, D., Buckner, R. L., Dale, A. M., Maguire, R. P., Hyman, B. T., Albert, M. S., & Killiany, R. J. (2006). An automated labeling system for subdividing the human cerebral cortex on MRI scans into gyral based regions of interest. NeuroImage, 31(3), 968–980. 10.1016/j.neuroimage.2006.01.021

13. Ding, N., Melloni, L., Zhang, H., Tian, X., & Poeppel, D. (2016). Cortical tracking of hierarchical linguistic structures in connected speech. Nature Neuroscience, 19(1), 158–164. 10.1038/nn.4186

14. Endress, A. D. (2024). Hebbian learning can explain rhythmic neural entrainment to statistical regularities. Developmental Science, 27(4), e13487. 10.1111/desc.13487

15. Evans, J. L., Saffran, J. R., & Robe-Torres, K. (2009). Statistical Learning in Children With Specific Language Impairment. Journal of Speech, Language, and Hearing Research, 52(2), 321–335. 10.1044/1092-4388(2009/07-0189)

16. Finestack, L. H., Richmond, E. K., & Abbeduto, L. (2009). Language Development in Individuals with Fragile X Syndrome. Topics in Language Disorders, 29(2), 133–148. 10.1097/tld.0b013e3181a72016

17. Fiser, J., & Aslin, R. N. (2001). Unsupervised Statistical Learning of Higher-Order Spatial Structures from Visual Scenes. Psychological Science, 12(6), 499–504. 10.1111/1467-9280.00392

18. Gabay, Y., Thiessen, E. D., & Holt, L. L. (2015). Impaired Statistical Learning in Developmental Dyslexia. *Journal of Speech*, Language, and Hearing Research: JSLHR, 58(3), 934–945. 10.1044/2015_JSLHR-L-14-0324

19. Gibson, J. R., Bartley, A. F., Hays, S. A., & Huber, K. M. (2008). Imbalance of Neocortical Excitation and Inhibition and Altered UP States Reflect Network Hyperexcitability in the Mouse Model of Fragile X Syndrome. Journal of Neurophysiology, 100(5), 2615–2626. 10.1152/jn.90752.2008

20. Giraud, A.-L., & Poeppel, D. (2012). Cortical oscillations and speech processing: Emerging computational principles and operations. Nature Neuroscience, 15(4), 511–517. 10.1038/nn.3063

21. Goel, A., Cantu, D. A., Guilfoyle, J., Chaudhari, G. R., Newadkar, A., Todisco, B., De Alba, D., Kourdougli, N., Schmitt, L. M., Pedapati, E., Erickson, C. A., & Portera-Cailliau, C. (2018). Impaired perceptual learning in a mouse model of Fragile X syndrome is mediated by parvalbumin neuron dysfunction and is reversible. Nature Neuroscience, 21(10), 1404– 1411. 10.1038/s41593-018-0231-0

22. Gramfort, A., Papadopoulo, T., Olivi, E., & Clerc, M. (2010). OpenMEEG: Opensource sosware for quasistatic bioelectromagnetics. BioMedical Engineering OnLine, 9(1), 45. 10.1186/1475-925X-9-45

23. Gross, J., Hoogenboom, N., Thut, G., Schyns, P., Panzeri, S., Belin, P., & Garrod, S. (2013). Speech Rhythms and Multiplexed Oscillatory Sensory Coding in the Human Brain. PLoS Biology, 11(12), e1001752. 10.1371/journal.pbio.1001752

24. Haebig, E., Saffran, J. R., & Weismer, S. E. (2017). Statistical Word Learning in Children with Autism Spectrum Disorder and Specific Language Impairment. Journal of Child Psychology and Psychiatry, and Allied Disciplines, 58(11), 1251–1263. 10.1111/jcpp.12734

25. Hagerman, R. J., Berry-Kravis, E., Hazlett, H. C., Bailey, D. B., Moine, H., Kooy, R. F., Tassone, F., Gantois, I., Sonenberg, N., Mandel, J. L., & Hagerman, P. J. (2017). Fragile X syndrome. Nature Reviews. Disease Primers, 3(PMID: 28960184), 17065. 10.1038/nrdp.2017.65

26. Henin, S., Turk-Browne, N. B., Friedman, D., Liu, A., Dugan, P., Flinker, A., Doyle, W., Devinsky, O., & Melloni, L. (2021). Learning hierarchical sequence representations across human cortex and hippocampus. Science Advances, 7(8), eabc4530. 10.1126/sciadv.abc4530

27. Herrera-Chaves, D., Gilmore, G., Abbass, M., Muller, L., Mirsattari, S., Kohler, S., & Batterink, L. (n.d.). The role of modality-specific brain regions in statistical learning: Insights from intracranial neural entrainment. Journal of Cognitive Neuroscience.

28. Hoffmann, A. (2022). Communication in fragile X syndrome: Patterns and implications for assessment and intervention. Frontiers in Psychology, 13(PMID: 36619013), 929379. 10.3389/fpsyg.2022.929379

29. Hu, A., Kozloff, V., Owen Van Horne, A., Chugani, D., & Qi, Z. (2024). Dissociation Between Linguistic and Nonlinguistic Statistical Learning in Children with Autism. Journal of Autism and Developmental Disorders, 54(5), 1912–1927. 10.1007/s10803-023-05902-1

30. Jeste, S. S., Kirkham, N., Senturk, D., Hasenstab, K., Sugar, C., Kupelian, C., Baker, E., Sanders, A. J., Shimizu, C., Norona, A., Paparella, T., Freeman, S. F. N., & Johnson, S. P. (2015). Electrophysiological evidence of heterogeneity in visual statistical learning in young children with ASD. Developmental Science, 18(1), 90–105. 10.1111/desc.12188

31. Jones, R. M., Tarpey, T., Hamo, A., Carberry, C., Brouwer, G., & Lord, C. (2018). Statistical Learning is Associated with Autism Symptoms and Verbal Abilities in Young Children with Autism. Journal of Autism and Developmental Disorders, 48(10), 3551–3561. 10.1007/s10803-018-3625-7

32. Karuza, E. A., Newport, E. L., Aslin, R. N., Starling, S. J., Tivarus, M. E., & Bavelier, D. (2013). The neural correlates of statistical learning in a word segmentation task: An fMRI study. Brain and Language, 127(1), 46–54. 10.1016/j.bandl.2012.11.007

33. Kidd, E. (2012). Implicit statistical learning is directly associated with the acquisition of syntax. Developmental Psychology, 48(1), 171–184. 10.1037/a0025405

34. Knox, A., Schneider, A., Abucayan, F., Hervey, C., Tran, C., Hessl, D., & Berry-Kravis, E. (2012). Feasibility, reliability, and clinical validity of the Test of Attentional Performance for Children (KiTAP) in Fragile X syndrome (FXS). Journal of Neurodevelopmental Disorders, 4(1), 2. 10.1186/1866-1955-4-2

35. Lammertink, I., Boersma, P., Wijnen, F., & Rispens, J. (2017). Statistical Learning in Specific Language Impairment: A Meta-Analysis. Journal of Speech, Language, and Hearing Research, 60(12), 3474–3486. 10.1044/2017_JSLHR-L-16-0439

36. Mayo, J., & Eigsti, I.-M. (2012). Brief report: A comparison of statistical learning in school-aged children with high functioning autism and typically developing peers. Journal of Autism and Developmental Disorders, 42(11), 2476–2485. 10.1007/s10803-012-1493-0

37. McNealy, K., Mazziotta, J. C., & Dapretto, M. (2006). Cracking the Language Code: Neural Mechanisms Underlying Speech Parsing. Journal of Neuroscience, 26(29), 7629–7639. 10.1523/JNEUROSCI.5501-05.2006

38. Moreau, C. N., Joanisse, M. F., Mulgrew, J., & Batterink, L. J. (2022). No statistical learning advantage in children over adults: Evidence from behaviour and neural entrainment. Developmental Cognitive Neuroscience, 57, 101154. 10.1016/j.dcn.2022.101154

39. Mourad, N., Reilly, J. P., de Bruin, H., Hasey, G., & MacCrimmon, D. (2007). A Simple and Fast Algorithm for Automatic Suppression of High-Amplitude Artifacts in EEG Data. 2007 IEEE International Conference on Acoustics, Speech and Signal Processing – ICASSP ‘07, 1, I-393–I-396. 10.1109/ICASSP.2007.366699

40. Nencheva, M. L., Peng, R., Tamir, D. I., & Lew-Williams, C. (2025). Infants track patterns of emoMon transiMons in the home. Journal of Experimental Psychology. Learning, Memory, and Cognition. 10.1037/xlm0001495

41. Neural Evidence for Linguistic Statistical Learning is Independent of Rhythmic and Cognitive Abilities in Neurotypical Adults: A Registered Report. (2025, June 16). Sciety. https://sciety-labs.elifesciences.org/articles/by?article_doi=10.31219/osf.io/qfsxk_v1

42. Obeid, R., Brooks, P. J., Powers, K. L., Gillespie-Lynch, K., & Lum, J. A. G. (2016). Statistical Learning in Specific Language Impairment and Autism Spectrum Disorder: A Meta-Analysis. Frontiers in Psychology, 7(PMID: 27602006), 1245. 10.3389/fpsyg.2016.01245

43. Obleser, J., & Kayser, C. (2019). Neural Entrainment and Attentional Selection in the Listening Brain. Trends in Cognitive Sciences, 23(11), 913–926. 10.1016/j.tics.2019.08.004

44. Pedapati, E. V., Ethridge, L. E., Liu, Y., Liu, R., Sweeney, J. A., DeStefano, L. A., Miyakoshi, M., Razak, K., Schmitt, L. M., Moore, D. R., Gilbert, D. L., Wu, S. W., Smith, E., Shaffer, R. C., Dominick, K. C., Horn, P. S., Binder, D., & Erickson, C. A. (2025). Frontal cortex hyperactivation and gamma desynchrony in Fragile X syndrome: Correlates of auditory hypersensitivity. PLOS One, 20(5), e0306157. 10.1371/journal.pone.0306157

45. Pedapati, E. V., Schmitt, L. M., Ethridge, L. E., Miyakoshi, M., Sweeney, J. A., Liu, R., Smith, E., Shaffer, R. C., Dominick, K. C., Gilbert, D. L., Wu, S. W., Horn, P. S., Binder, D. K., Lamy, M., Axford, M., & Erickson, C. A. (2022). Neocortical localization and thalamocortical modulation of neuronal hyperexcitability contribute to Fragile X Syndrome. Communications Biology, 5(1), 442. 10.1038/s42003-022-03395-9

46. Poeppel, D., & Assaneo, M. F. (2020). Speech rhythms and their neural foundations. Nature Reviews Neuroscience, 21(6), 322–334. 10.1038/s41583-020-0304-4

47. Raviv, L., & Arnon, I. (2018). The developmental trajectory of children’s auditory and visual statistical learning abilities: Modality-based differences in the effect of age. Developmental Science, 21(4), e12593. 10.1111/desc.12593

48. Reiss, A. L., Lee, J., & Freund, L. (1994). Neuroanatomy of fragile X syndrome: The temporal lobe. Neurology, 44(7), 1317–1324. 10.1212/wnl.44.7.1317

49. Richter, C. G., Thompson, W. H., Bosman, C. A., & Fries, P. (2015). A jackknife approach to quantifying single-trial correlation between covariance-based metrics undefined on a single-trial basis. NeuroImage, 114, 57–70. 10.1016/j.neuroimage.2015.04.040

50. Romberg, A. R., & Saffran, J. R. (2010). Statistical language learning: Mechanisms and constraints. Current Directions in Psychological Science, 19(2), 114–118. 10.1177/0963721410365001

51. Saffran, J. R., Aslin, R. N., & Newport, E. L. (1996). Statistical Learning by 8-Month-Old Infants. *Science*, New Series, 274(5294), 1926–1928.

52. Saffran, J. R., & Kirkham, N. Z. (2018). Infant Statistical Learning. Annual Review of Psychology, 69(1), 181–203. 10.1146/annurev-psych-122216-011805

53. Saffran, J. R., Newport, E. L., & Aslin, R. N. (1996). Word Segmentation: The Role of Distributional Cues. Journal of Memory and Language, 35(4), 606–621. 10.1006/jmla.1996.0032

54. Saffran, J. R., Newport, E. L., Aslin, R. N., Tunick, R. A., & Barrueco, S. (1997). Incidental Language Learning: Listening (and Learning) Out of the Corner of Your Ear. Psychological Science, 8(2), 101–105. 10.1111/j.1467-9280.1997.tb00690.x

55. Sahin, M., Jones, S. R., Sweeney, J. A., Berry-Kravis, E., Connors, B. W., Ewen, J. B., Hartman, A. L., Levin, A. R., Potter, W. Z., & Mamounas, L. A. (2019). Discovering translational biomarkers in neurodevelopmental disorders. Nature Reviews Drug Discovery, 18(4), 235–236. 10.1038/d41573-018-00010-7

56. Sandoval, G. M., Shim, S., Hong, D. S., Garrett, A. S., QuinMn, E.-M., Marzelli, M. J., Patnaik, S., Lightbody, A. A., & Reiss, A. L. (2018). Neuroanatomical abnormalities in fragile X syndrome during the adolescent and young adult years. Journal of Psychiatric Research, 107, 138–144. 10.1016/j.jpsychires.2018.10.014

57. Schmitt, L. M., Arzuaga, A. L., Dapore, A., Duncan, J., Patel, M., Larson, J. R., Erickson, C. A., Sweeney, J. A., & Ragozzino, M. E. (2023). Parallel learning and cognitive flexibility impairments between Fmr1 knockout mice and individuals with fragile X syndrome. Frontiers in Behavioral Neuroscience, 16, 1074682. 10.3389/fnbeh.2022.1074682

58. Schmitt, L. M., Li, J., Liu, R., Horn, P. S., Sweeney, J. A., Erickson, C. A., & PedapaM, E. V. (2022). Altered frontal connectivity as a mechanism for executive function deficits in fragile X syndrome. Molecular Autism, 13(1), 47. 10.1186/s13229-022-00527-0

59. Scott-Van Zeeland, A. A., McNealy, K., Wang, A. T., Sigman, M., Bookheimer, S. Y., & Dapretto, M. (2010). No Neural Evidence of Statistical Learning During Exposure to Artificial Languages in Children with Autism Spectrum Disorders. Biological Psychiatry, 68(4), 345–351. 10.1016/j.biopsych.2010.01.011

60. Sherman, B. E., Graves, K. N., & Turk-Browne, N. B. (2020). The prevalence and importance of statistical learning in human cognition and behavior. Current Opinion in Behavioral Sciences, 32, 15–20. 10.1016/j.cobeha.2020.01.015

61. Siegelman, N., & Frost, R. (2015). Statistical learning as an individual ability: Theoretical perspectives and empirical evidence. Journal of Memory and Language, 81, 105–120. 10.1016/j.jml.2015.02.001

62. Smalle, E. H. M., Daikoku, T., Szmalec, A., Duyck, W., & Möttönen, R. (2022). Unlocking adults’ implicit statistical learning by cognitive depletion. Proceedings of the National Academy of Sciences, 119(2), e2026011119. 10.1073/pnas.2026011119

63. Theeuwes, J., Huang, C., Frings, C., & van Moorselaar, D. (2024). Statistical learning of motor preparation. Journal of Experimental Psychology. Human Perception and Performance, 50(2), 152–162. 10.1037/xhp0001174

64. Van Der Molen, M. J. W., Stam, C. J., & Van Der Molen, M. W. (2014). Resting-State EEG Oscillatory Dynamics in Fragile X Syndrome: Abnormal Functional Connectivity and Brain Network Organization. PLoS ONE, 9(2), e88451. 10.1371/journal.pone.0088451

65. Van der Molen, M. J. W., & Van der Molen, M. W. (2013). Reduced alpha and exaggerated theta power during the resting-state EEG in fragile X syndrome. Biological Psychology, 92(2), 216–219. 10.1016/j.biopsycho.2012.11.013

66. Vandermosten, M., Wouters, J., Ghesquière, P., & Golestani, N. (2019). Statistical Learning of Speech Sounds in Dyslexic and Typical Reading Children. Scientific Studies of Reading, 23(1), 116–127. 10.1080/10888438.2018.1473404

67. Wagley, N., Lajiness-O’Neill, R., Hay, J. S. F., Ugolini, M., Bowyer, S. M., Kovelman, I., & Brennan, J. R. (2020). Predictive Processing during a Naturalistic Statistical Learning Task in ASD. eNeuro, 7(6). 10.1523/ENEURO.0069-19.2020

68. Wang, J., Ethridge, L. E., Mosconi, M. W., White, S. P., Binder, D. K., PedapaM, E. V., Erickson, C. A., Byerly, M. J., & Sweeney, J. A. (2017). A resting EEG study of neocortical hyperexcitability and altered functional connectivity in fragile X syndrome. Journal of Neurodevelopmental Disorders, 9(PMID: 28316753), 11. 10.1186/s11689-017-9191-z

69. Wang, X., Sela-Donenfeld, D., & Wang, Y. (2023). Axonal and presynaptic FMRP: Localization, signal, and functional implications. Hearing Research, 430, 108720. 10.1016/j.heares.2023.108720

70. Waschke, L., Tune, S., & Obleser, J. (2019). Local cortical desynchronization and pupil-linked arousal differentially shape brain states for optimal sensory performance. eLife, 8, e51501. 10.7554/eLife.51501

71. Yi, H. G., Leonard, M. K., & Chang, E. F. (2019). The Encoding of Speech Sounds in the Superior Temporal Gyrus. Neuron, 102(6), 1096–1110. 10.1016/j.neuron.2019.04.023

72. Zhang, M., Riecke, L., & Bonte, M. (2021). Neurophysiological tracking of speech-structure learning in typical and dyslexic readers. Neuropsychologia, 158, 107889. 10.1016/j.neuropsychologia.2021.107889

73. Zioga, I., Weissbart, H., Lewis, A. G., Haegens, S., & Martin, A. E. (2023). Naturalistic spoken language comprehension is supported by alpha and beta oscillations. The Journal of Neuroscience, JN-RM-1500-22. 10.1523/JNEUROSCI.1500-22.2023

