## Supplemental Table 1 for "Aberrant Neural Entrainment to Word-Level Speech Patterns in Fragile X Syndrome: Evidence for a Statistical Learning Deficit"

|  | **ZITC_Word** | | |
| --- | --- | --- | --- |
| *Predictors* | *Estimates* | *CI* | *p* |
| (Intercept) | 0.27 | 0.15 – 0.39 | **<0.001** |
| CoreRegion [PF] | -0.15 | -0.31 – 0.01 | 0.061 |
| CoreRegion [F] | -0.08 | -0.23 – 0.06 | 0.254 |
| CoreRegion [T] | 0.40 | 0.29 – 0.51 | **<0.001** |
| CoreRegion [C] | -0.05 | -0.23 – 0.13 | 0.572 |
| CoreRegion [L] | -0.03 | -0.19 – 0.13 | 0.712 |
| CoreRegion [P] | 0.02 | -0.14 – 0.18 | 0.797 |
| Hemisphere [L] | 0.00 | -0.06 – 0.06 | 0.994 |
| Group [FXS] | -0.08 | -0.27 – 0.11 | 0.403 |
| CoreRegion [PF] × Hemisphere [L] | 0.06 | -0.10 – 0.21 | 0.479 |
| CoreRegion [F] × Hemisphere [L] | -0.01 | -0.16 – 0.13 | 0.855 |
| CoreRegion [T] × Hemisphere [L] | -0.05 | -0.16 – 0.06 | 0.356 |
| CoreRegion [C] × Hemisphere [L] | 0.02 | -0.16 – 0.19 | 0.853 |
| CoreRegion [L] × Hemisphere [L] | -0.01 | -0.17 – 0.15 | 0.897 |
| CoreRegion [P] × Hemisphere [L] | -0.06 | -0.21 – 0.10 | 0.472 |
| CoreRegion [PF] × Group [FXS] | -0.11 | -0.33 – 0.11 | 0.323 |
| CoreRegion [F] × Group [FXS] | 0.09 | -0.11 – 0.29 | 0.361 |
| CoreRegion [T] × Group [FXS] | -0.25 | -0.40 – -0.09 | **0.002** |
| CoreRegion [C] × Group [FXS] | 0.19 | -0.06 – 0.44 | 0.134 |
| CoreRegion [L] × Group [FXS] | 0.05 | -0.17 – 0.27 | 0.632 |
| CoreRegion [P] × Group [FXS] | -0.00 | -0.22 – 0.22 | 0.996 |
| Hemisphere [L] × Group [FXS] | 0.04 | -0.05 – 0.13 | 0.367 |
| (CoreRegion [PF] × Hemisphere [L]) × Group [FXS] | -0.06 | -0.28 – 0.16 | 0.591 |
| (CoreRegion [F] × Hemisphere [L]) × Group [FXS] | -0.08 | -0.27 – 0.12 | 0.461 |
| (CoreRegion [T] × Hemisphere [L]) × Group [FXS] | 0.07 | -0.09 – 0.22 | 0.389 |
| (CoreRegion [C] × Hemisphere [L]) × Group [FXS] | 0.15 | -0.10 – 0.40 | 0.242 |
| (CoreRegion [L] × Hemisphere [L]) × Group [FXS] | -0.07 | -0.29 – 0.15 | 0.550 |
| (CoreRegion [P] × Hemisphere [L]) × Group [FXS] | 0.09 | -0.13 – 0.31 | 0.411 |
| **Random Effects** | | | |
| σ^2^ | 1.29 | | |
| τ_00_ _Node_ | 0.02 | | |
| τ_00_ _SubjectID_ | 0.08 | | |
| ICC | 0.07 | | |
| N _SubjectID_ | 48 | | |
| N _Node_ | 68 | | |
| Observations | 3264 | | |
| Marginal R^2^ / Conditional R^2^ | 0.031 / 0.100 | | |
